# Automatic instance segmentation of mitochondria in electron microscopy data

**DOI:** 10.1101/2021.05.24.444785

**Authors:** Luke Nightingale, Joost de Folter, Helen Spiers, Amy Strange, Lucy M Collinson, Martin L Jones

**Affiliations:** The Francis Crick Institute, London, NW1 1AT, United Kingdom; University of Oxford, Oxford, OX1 3RH, United Kingdom

**Keywords:** Electron microscopy, Mitochondria, Instance Segmentation, Deep Learning, 3D

## Abstract

We present a new method for rapid, automated, large-scale 3D mitochondria instance segmentation, developed in response to the ISBI 2021 MitoEM Challenge. In brief, we trained separate machine learning algorithms to predict (1) mitochondria areas and (2) mitochondria boundaries in image volumes acquired from both rat and human cortex with multi-beam scanning electron microscopy. The predictions from these algorithms were combined in a multi-step post-processing procedure, that resulted in high semantic and instance segmentation performance. All code is provided via a public repository.

## 1. INTRODUCTION

Electron microscopy has undergone a radical advancement in data generation capacity through the development of a series of automated systems that are able to rapidly collect thousands of serial images through whole cell and tissues. This has displaced the research bottleneck in this domain to data analysis – one of the greatest challenges faced by modern imaging is how to efficiently and quickly extract meaningful insight from the incredibly rich data produced.

However, the development of novel analytical pipelines is often hampered by a scarcity of relevant, publicly available, well-labelled datasets. In response, the MitoEM dataset, two 3D mitochondria instance segmentations derived from rat and human cortex volume EM image stacks, were created and shared [1] along with a challenge to the community - to develop novel robust automated instance segmentation approaches for this data. Here, we present our response to this challenge.

## 2. PROPOSED METHOD

### 2.1. Overview of approach

Following acquisition and pre-processing of challenge data [1] (Sections 2.2, 2.3), we trained a series of customised 3D U-Nets (Sections 2.4, 2.6, 2.7) to predict mitochondria areas and boundaries (Section 2.8) in both the rat and human data. The predictions from these algorithms were combined in a multi-step post-processing procedure (Section 2.9), that resulted in high semantic and instance segmentation performance (Table 1). An overview of the approach used is presented in Figure 1.

**Fig. 1.**
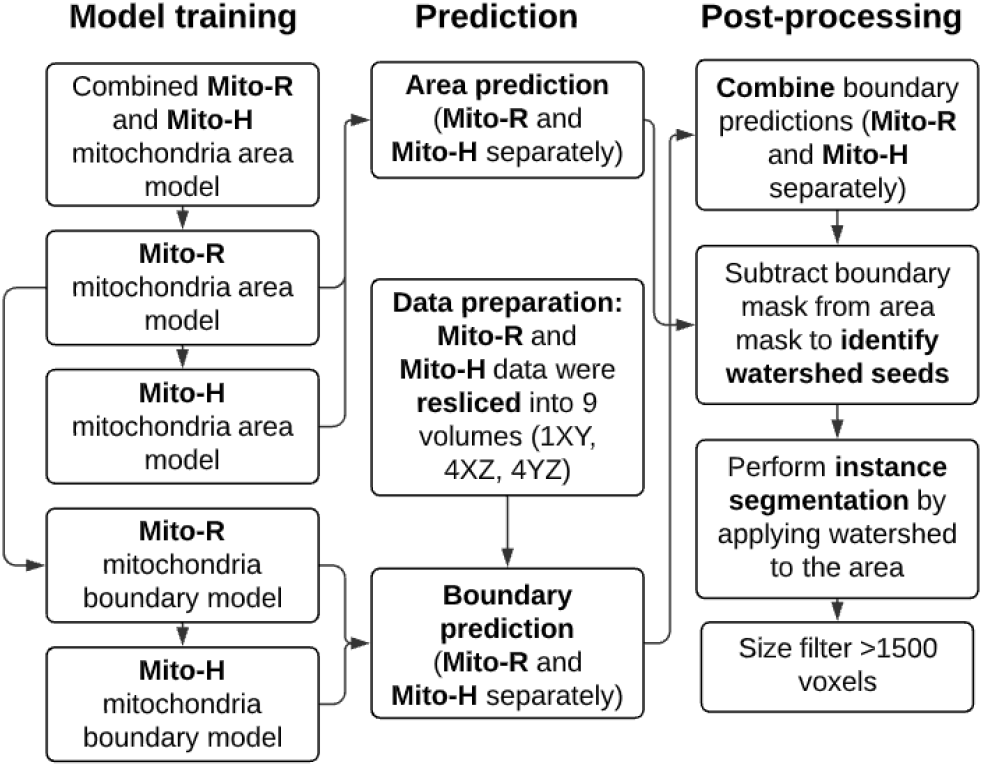
Analytical pipeline: model training, prediction and post-processing

**Table 1.** Model performance metrics, computed on the unseen validation data. Semantic segmentation metrics are reported for single-axis (XY) prediction.

|  | Dice | IoU | Precision | Recall |
| --- | --- | --- | --- | --- |
| Mito-R area | 0.946 | 0.897 | 0.959 | 0.933 |
| Mito-R boundary | 0.835 | 0.719 | 0.810 | 0.862 |
| Mito-H area | 0.920 | 0.852 | 0.934 | 0.906 |
| Mito-H boundary | 0.829 | 0.707 | 0.817 | 0.841 |

|  | AP@75 | AP@75<br>large | AP@75<br>medium | AP@75<br>small |
| --- | --- | --- | --- | --- |
| Rat | 0.715 | 0.787 | 0.404 | 0.007 |
| Human | 0.625 | 0.734 | 0.478 | 0.034 |

### 2.2. Data acquisition and annotation

Two datasets were provided for the development of 3D mitochondria instance segmentation approaches in association with the MitoEM Challenge [1]. The two 30 µm^3^ image volumes provided were acquired from rat (Mito-R) and human cortex (Mito-H) (1000*×* 4096 *×*4096 in voxels at 30*×* 8*×* 8 nm resolution) with multi-beam scanning electron microscopy (for a full description of data acquisition see [1]).

For both volumes, 1000 consecutive slices were provided. Ground truth instance segmentation was generated for the two volumes through an iterative, semi-automated approach, that applied a 3D U-Net in collaboration with manual correction by expert annotators [1]. Ground truth mitochondria instance labels were provided for the first 500 slices, with the first 400 (0–399) used for training, and the remaining (400–499) used for validation.

### 2.3. Pre-processing of training data

For both the Mito-R and Mito-H datasets, the 400 training slices were each partitioned into four sets of 100. Each 100 slice stack was further divided into four equal tiled stacks, resulting in a total of 16 stacks per dataset for training. The 100 slices of validation data were similarly split into four tiled stacks for each dataset and used for validation error evaluation during training.

Ground truth data for the mitochondria instance boundaries were generated using OpenCV’s `findContours` function [2] on a per-instance basis for each 2D plane per stack. The boundaries were set at a width of 5 pixels in approximate correspondence with the membrane thickness in the raw data.

### 2.4. Model Architecture

Using the mitochondrial data provided in the challenge and the derived boundary ground truth data (Section 2.3) four separate CNNs were trained to provide semantic segmentation predictions of the (1) Mito-R mitochondria areas (2) Mito-H mitochondria areas (3) Mito-R mitochondria boundaries (4) Mito-H mitochondria boundaries.

All CNNs used the same custom 3D U-Net architecture. The individual blocks were inspired by the Inception CNN [3], having the same pattern of convolutions, pooling and activation, as well as skip connections within the block. Overall network topology was similar to the U-Net [4] - an encoder path of four blocks and decoder path of four blocks with skip connections between them. To build on our former work establishing a CNN for prediction of the nuclear envelope [5], we selected an input patch size consistent with this previous model (12, 256, 256). This enabled us to initialize our new model using the nuclear envelope weights.

Additional model parameters were as follows; dropout rate of 0.3, 32 start filters, Adam optimizer with an initial learning rate of 0.0001. The model was trained using an *AWS EC2 P3*.*2xlarge* instance. The loss function used for the model was the smoothed dice coefficient (or F-measure), where

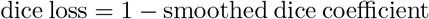

### 2.5. Model performance metrics

In line with the MitoEM challenge [1], we report the average precision (AP-75) metric for instance segmentation, requiring at least 75% intersection over union (IoU) with ground-truth for a true positive detection. For completeness, we also report the Dice coefficient and IoU as a measure of the semantic segmentation performance of our model.

### 2.6. Model training – mitochondria areas

First, a model was trained with combined Mito-R and Mito-H mitochondria area data. This model was initialized with the weights established in prior work for the semantic segmentation of the nuclear envelope [5] (Section 2.4). We trained for 100 epochs on the combined Mito-R and Mito-H data, with each epoch comprising 100 iterations of the Adam optimizer and batches of 12 samples. Mito-R and Mito-H data were presented alternately, for both training and validation, hence a Mito-R validation stack was occasionally presented after a Mito-H epoch (and vice versa).

The weights from this combined Mito-R and Mito-H mitochondria area model were used for the initialization of a Mito-R mitochondria area model. This model was trained for 183 epochs (stopping after convergence). Due to memory constraints, one of the 16 (Section 2.3) training stacks was presented per epoch. This resulted in each individual stack appearing once per 16 epochs. To check for convergence, we waited until most individual stacks had approximately constant training error from one complete cycle to the next. We selected the training epoch in the last complete cycle that had the best performance as the final model. Applying the final Mito-R mitochondria area model on the full rat validation dataset showed a high semantic segmentation performance (Dice = 0.946; IoU = 0.897, Table 1).

Next, the weights from the Mito-R mitochondria area model were used for the initialization of a Mito-H mitochondria area model. This model was trained for 526 epochs (stopping after convergence), and again one of the 16 (Section 2.3) training stacks was presented per epoch. Applying the final Mito-H mitochondria area model on the full human validation dataset showed a high semantic segmentation performance (Dice = 0.920; IoU = 0.852, Table 1).

### 2.7. Model training – mitochondria boundaries

Following training of the two mitochondria area models (Section 2.6), we next trained separate models to identify mitochondria boundaries in the rat and human data. First, the weights from the Mito-R mitochondria area model were used to initialize the Mito-R mitochondria boundary model. This model was trained for 812 epochs, and, as with the area models, one of the 16 training stacks was presented per epoch. Applying the Mito-R mitochondria boundary model on the full rat validation achieved a Dice = 0.835 and IoU = 0.717 (Table 1).

Finally, the weights from the Mito-R mitochondria boundary model were used to initialize the Mito-H mitochondria boundary model. This model was trained for 665 epochs, and, as with the area models, one of the 16 training stacks was presented per epoch. Applying the Mito-H mitochondria boundary model on the full human validation achieved a Dice = 0.829 and IoU = 0.707 (Table 1).

### 2.8. Improving predicted mitochondrial boundaries with Tri-Axis Prediction

The prediction of boundary segmentations can often fail in regions where the boundary is oriented parallel to the imaging plane due to a lack of distinct morphological features and poor contrast. Hence, for a more robust mitochondrial boundary prediction, here we build upon the *Tri-Axis Prediction* (TAP) method developed in [5] (Figure 2).

**Fig. 2.**
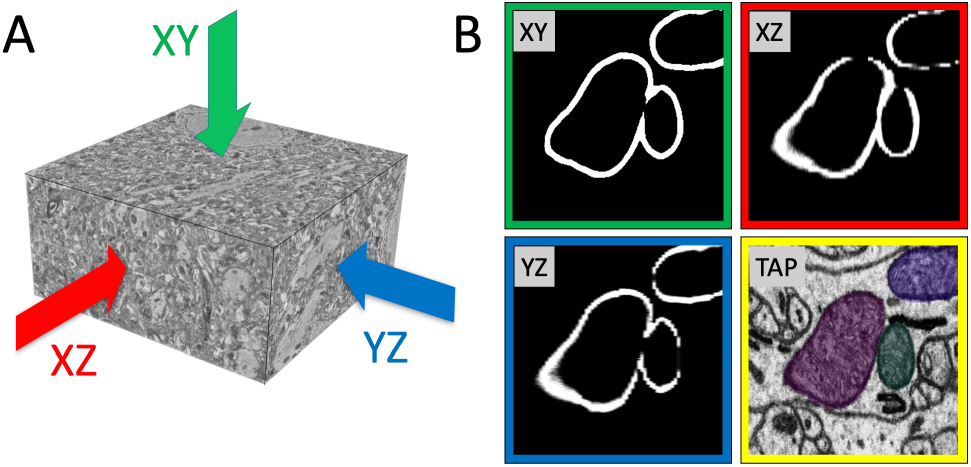
Tri-Axis Prediction (TAP) schematic for Mito-R boundary prediction data (A). Multi-view predictions on a cropped region (B). Boundary gaps in the original XY prediction cause erroneous joining of watershed seeds. By combining predictions from all three views with TAP, boundary integrity is more robustly maintained, giving better watershed seed identification for instance segmentation (yellow border).

Additional mitochondria boundary predictions were generated for orthogonal axes by reslicing the data in the XZ and YZ planes. To approximately match the training data voxel size after rotation, the voxel z-scale was interpolated from 30 nm to 10 nm using cubic interpolation in Fiji [6], resulting in a stack with a voxel size of 8 *×* 8 *×* 10 nm and 3*×* more z-slices.

To approximately match the voxel size used for boundary model training, and to avoid further downscaling interpolation of the raw data, stacks with a slice separation of 32 nm were created by deinterleaving the rescaled stack into four stacks with XYZ voxel size of 8*×* 32 *×* 10 nm for the XZ view and 32*×* 8 *×* 10 nm for the YZ view.

The mitochondria boundary prediction model was then run over each deinterleaved stack for each view, resulting in a total of 9 prediction masks (one from the original XY orientation and four each from the XZ and YZ views). The prediction stacks were then transformed back to the original orientation by rotation and interpolation. The final boundary mask was generated using a simple voting method where one or more votes designated a voxel as part of a boundary.

### 2.9. Identification of mitochondrial instances

The generation of both mitochondrial area and boundary predictions enabled application of a watershed-based method [1] to generate mitochondrial instance segmentations (Figure 1). To create seed areas for the marker-based watershed, the boundary mask was subtracted from the area mask to produce disconnected 3D ‘islands’. This method also facilitated the removal of small objects. Running the watershed algorithm from these seeds allowed separation of the semantic segmentation into individual instances. Quantitative (Table 1) and qualitative inspection (Figure 3, Supplementary Movies 1–6) of the results of this approach indicate good performance.

**Fig. 3.**
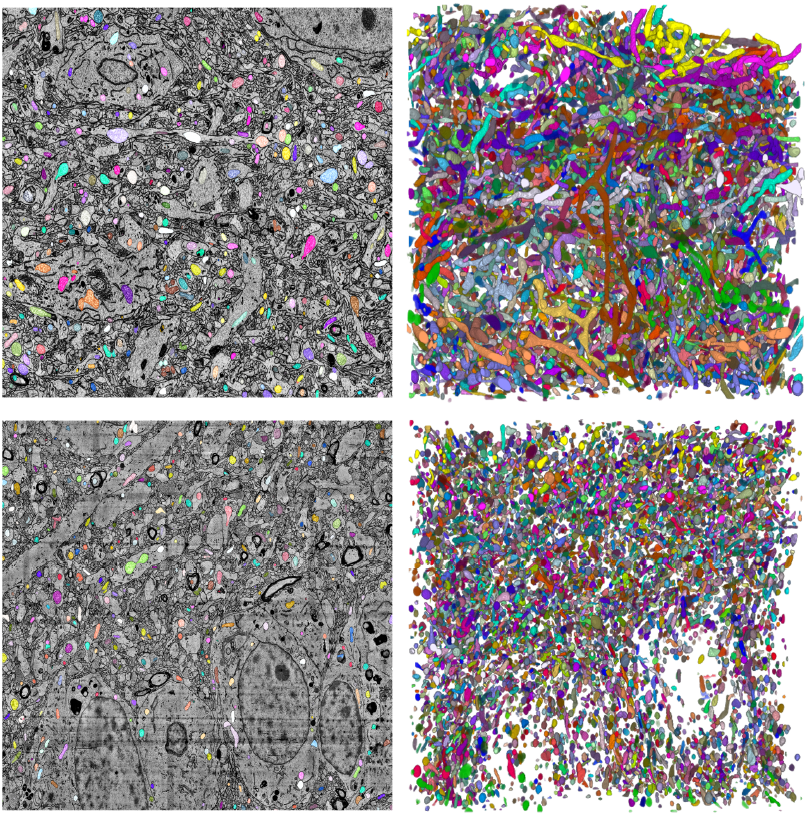
Performance of instance segmentation approach in Mito-R data (top left) and Mito-H data (bottom left), (shown for slice 250 of 500) shown alongside a 3D rendering of all segmented mitochondrial instances for each full volume.

### 2.10. Computing Resource and code availability

Data analysis was performed on available high performance computing resources. This included Amazon Web Services (AWS) cloud computing service (www.aws.amazon.com), and a local high performance compute cluster called “CAMP” (Crick Data Analysis and Management Platform). Code is available at: https://github.com/FrancisCrickInstitute/mitoem-challenge.

## 3. DISCUSSION AND CONCLUSION

We present here our response to the ISBI 2021 MitoEM Challenge, in which we produce mitochondrial instance segmentations for two image volumes acquired from rat and human cortex with multi-beam scanning electron microscopy. In summary, our approach involved the training of multiple machine learning algorithms for prediction of mitochondria areas and boundaries, which were subsequently used in conjunction with a series of post-processing steps to determine mitochondrial instances. Through quantitative performance metrics and qualitative inspection of the data produced, our proposed approach was found to be robust. We note that the semantic segmentation metrics we present show much higher performance than our instance segmentation, suggesting there is scope to improve the performance of our proposed approach, through mechanisms such as alternative model training strategies.

## 4. COMPLIANCE WITH ETHICAL STANDARDS

This is a numerical simulation study for which no ethical approval was required.

## 5. ACKNOWLEDGMENTS

This work was supported by the Francis Crick Institute which receives its core funding from Cancer Research UK (FC001999), the UK Medical Research Council (FC001999), and the Wellcome Trust (FC001999). The authors declare no competing interests. For the purpose of Open Access, the author has applied a CC BY public copyright licence to any Author Accepted Manuscript version arising from this submission

## 7. SUPPLEMENTARY MATERIAL

**Supplementary Movie 1:** Mito-R data with overlaid area prediction (magenta) and boundary prediction (green)

**Supplementary Movie 2:** Mito-H data with overlaid area prediction (magenta) and boundary prediction (green)

**Supplementary Movie 3:** Mito-R data with overlaid instance segmentation

**Supplementary Movie 4:** Mito-H data with overlaid instance segmentation

**Supplementary Movie 5:** 3D rendering of Mito-R instance segmentation

**Supplementary Movie 6:** 3D rendering of Mito-H instance segmentation

